# Quantitative Time-Course Analysis of Osmotic and Salt Stress in *Arabidopsis thaliana* using Short Gradient Multi-CV FAIMSpro BoxCar DIA

**DOI:** 10.1101/2023.02.22.529555

**Authors:** M.C. Rodriguez Gallo, Q. Li, M. Talasila, RG Uhrig

## Abstract

A major limitation when undertaking quantitative proteomic time-course experimentation is the tradeoff between depth-of-analysis and speed-of-analysis. In high complexity and high dynamic range sample types, such as plant extracts, balance between resolution and time is especially apparent. To address this, we evaluate multiple composition voltage (CV) High <u>F</u>ield <u>A</u>symetric Waveform <u>I</u>on <u>M</u>obility <u>S</u>pectrometry (FAIMSpro) settings using the latest label-free single-shot Orbitrap-based DIA acquisition workflows for their ability to deeply-quantify the *Arabidopsis thaliana* seedling proteome. Using a BoxCarDIA acquisition workflow with a −30 −50 −70 CV FAIMSpro setting we are able to consistently quantify >5000 *Arabidopsis* seedling proteins over a 21-minute gradient, facilitating the analysis of ~42 samples per day. Utilizing this acquisition approach, we then quantified proteome-level changes occurring in *Arabidopsis* seedling shoots and roots over 24 h of salt and osmotic stress, to identify early and late stress response proteins and reveal stress response overlaps. Here, we successfully quantify >6400 shoot and >8500 root protein groups, respectively, quantifying nearly ~9700 unique protein groups in total across the study. Collectively, we pioneer a short gradient, multi-CV FAIMSpro BoxCarDIA acquisition workflow that represents an exciting new analysis approach for undertaking quantitative proteomic time-course experimentation in plants.

## Introduction

The ability to perform quantitative proteomic time-course experimentation is critical to revealing dynamic proteome-level changes occurring in the molecular landscape of cells. This is particularly important in defining how plants perform cellular tasks throughout the day, execute development programs, and respond to stress. However, plant samples are high dynamic range, high complexity samples, creating a constant trade-off between speed-of-analysis and depth-of-analysis, with plant samples typically requiring longer chromatographic gradients, and therefore longer acquisition times to generate quantitative proteomic data (1, 2). This then restricts the ability to perform time-course or time-series proteomic experiments, as deeper proteome quantification comes with substantial acquisition-time overheads. However, the development of High <u>F</u>ield <u>A</u>symmetric Waveform <u>I</u>on <u>M</u>obility <u>S</u>pectrometry (FAIMSpro) has created new opportunities by decreasing competing precursor ion noise in mass spectrometry (MS) analyses. When coupled with microliter/min flow rates, FAIMSpro has been shown to be an effective tool for increasing the number of samples analyzed per day at comparable depths of analysis to longer gradients (3).

The majority of quantitative plant proteomics data continues to be generated using data dependent acquisition (DDA), which is known to increase missing values (MV) between replicate samples due to its “TopN” precursor selection approach (4, 5). Without additional offline sample processing, such as sample fractionation to reduce the complexity of injected samples, the semi-stochastic MS^1^ scan of DDA acquisitions will select more intense precursor ions for MS^2^ fragmentation, thus biasing analyses. The MV problem and the increased variability between DDA runs is especially problematic for large-scale time-course analyses. To address the MV problem and increase reproducibility between samples, data independent acquisition (DIA) methodologies can be applied. DIA removes the TopN precursor ion selection step based on MS^1^ intensity by selecting precursor ions across a mass range, followed by fragmentation and MS^2^ analysis (6).

Further gains in depth of analysis and reduction of MV have been achieved using sequential window MS^1^ acquisitions, such as BoxCarDIA, which segment precursor ions prior to MS^2^ analysis (5, 7, 8). Ultimately, MS methods dependent on *m/z* window fragmentation rather than peak intensity come at the expense of multiplexed fragment ion spectra that requires computational heavy identification and quantification algorithms in order to effectively match spectra to peptides. Historically, DIA spectral matching has relied on DDA-derived spectral libraries, increasing instrumentation run time, however, recent development of library-free methods such as, BoxCarDIA and direct DIA (dDIA), have capitalized on software that allow for the effective analysis of DIA data without the creation of a spectral library. Software capable of processing library-free DIA includes: commercially available products like Spectronaut (Biognosys AG; https://biognosys.com/) or freely available solutions like DIA-NN (9) or MaxDIA (10).

In addition to developing more efficient acquisition methods, orbitrap instrumentation has been shown to benefit from the use of FAIMSpro (3), whereby the compensation voltage (CV) is modulated to allow desired precursor ions to traverse through the FAIMSpro into the mass spectrometer (11). Implementation of FAIMSpro for proteome analyses has been undertaken with single-CV acquisitions, which has led to increases in protein identification (12, 13). Although single-CV FAIMSpro acquisitions improved protein identification, each CV will possess bias towards a sub-selection of precursor ions. Correspondingly, multi-CV acquisitions using separate injections of the same sample at different CVs has been explored to overcome the bias of a single-CV, however, this extends required MS acquisition time and requires increased sample amounts. As a result, researchers developed “CV stepping” which cycles multiple CVs in a single injection, thereby retaining the benefits of FAIMSpro without the need for multiple single-CV injections of the same sample (14).

Despite the detection improvements offered by FAIMSpro facilitated acquisition methods, pervasive limitations still exist for experiments consisting of hundreds of samples due to the use of nanoflow LC gradients. As such, FAIMSpro coupled with microflow LC gradients (e.g. >1 μl/min) could reduce run time while maintaining high peptide identifications. For example, Bekker-Jensen et al. (2020) reported improved peptide identification through the combined implementation of single-CV FAIMSpro with DIA and short LC gradients, enabling the processing of hundreds of mammalian samples in days rather than weeks. However, CV-stepping in conjunction with DIA and short LC gradients has not yet been explored. Therefore, we evaluated multi-CV FAIMSpro DIA for its ability to maintain depth of analysis when deployed with microflow LC gradients to analyze high dynamic range and high complexity plant samples.

For plant biology to fully elucidate molecular mechanisms, the ability to readily track changes in gene expression and/or protein abundance across time is required. Gene expression profiling technologies, such as RNAseq / microarrays, have led the way, being utilized to track transcriptional changes across time in response to environmental stress (15–17) and photoperiod (18, 19), amongst other conditions (20). These transcriptional time-course experiments have created invaluable repositories for targeted hypothesis development (e.g. Diurnal DB; http://diurnal.mocklerlab.org/). However, the ability to systematically and quantitatively track changes in protein abundance across time has lagged behind, with research efforts profiling proteome-level changes over time focusing on circadian / diel (2, 21, 22) and select stress (23–26) related questions. A major reason for the lack of time-course proteomic experimentation in plants is that comprehensive time-course proteome analyses require extensive MS run time, making these analyses time-consuming and costly. Consequently, MS methods geared at reducing run time, while maintaining high quality protein identification and quantification, are required.

Increased drought and salt stress on agricultural lands around the world as a result of climate change continues to reduce crop yield and economic gains (27). Owing to its accessibility and depth of analysis, transcriptome technologies have typically been employed as the primary research tool for identifying new breeding / biotechnology targets related to abiotic stress (28, 29). However, transcriptional changes often weakly correlate with protein-level changes (30), creating a need for time-course based proteomic analyses in order to fully understand how plants response to environmental stresses. By comparison to the abundance of transcriptome data, knowledge of abiotic stress responses at the protein-level in plants remains sparse. Therefore, it is critical to define plant osmotic and salt stress responses at the protein-level through time-course experimentation. To effectively do this at the protein-level, we developed a fast-flow, short-gradient, FAIMSpro enabled BoxCarDIA acquisition approach that utilizes CV stepping to rapidly generate a deep quantitative proteomic profile of plants. Our application of this acquisition approach allowed us to reliably elucidate a time-resolved understanding of how *Arabidopsis* seedling shoots and roots respond to osmotic and salt stress.

## Experimental Procedure

### Plant growth & Abiotic stress treatment

Sterilized *Arabidopsis thaliana* wild-type Col-0 (*Arabidopsis*) seeds were plated on 0.5 x MS (Caisson Laboratories Inc. Murashige & Skoog MSP01) 0.8 % plant agar (Caisson laboratories Inc. Phytoblend™ PTP01) and stratified for 4 d at 4 °C in the dark. Seedlings were germinated under a 16 h : 8 h photoperiod for a total 15 d (14 d before stress + 1 d of stress) with 100 μmol/m^2^/s LED lights at constant 22 °C. Seedlings were grown vertically in custom 3D printed blacked-out vertical plate holders to minimize root exposure to light. At ZT3 of 14 d, seedlings were transferred to either 150 mM NaCl, 300 mM mannitol, or 0.5 X MS control plates. Plants were harvested at 0 h, 1 h, 3 h, 6 h, 12 h, and 24 h post stress exposure.

### Tissue processing, protein extraction, and protein digest

Frozen plant material was ground using Geno/Grinder (SPEX SamplePrep) for 30 seconds at 1200 rpm and aliquoted into 100 mg (shoot) and 50 mg (root) fractions using liquid N_2_ to prevent tissue degradation. Ground tissue was then re-suspended in protein extraction buffer (50 mM HEPES-KOH pH 8.0, 100 mM NaCl, and 4 % (w/v) SDS) at a 1:2 (w/v) ratio and extracted by shaking 1000 RPM 95°C for 5 minutes using a tabletop shaker (Eppenforf ThermoMixer F2.0). Samples were then centrifuged at 20,000 x g for 10 minutes at room temperature then the supernatant was retained in new tubes. Protein extracts were then reduced with 10 mM dithiothreitol (D9779, Sigma), followed by alkylation with 30 mM Iodoacetamide (I1149, Sigma) for 30 minutes at room temperature. Total proteome peptide fractions were then generated using a KingFisher APEX (Thermo Scientific) automated sample processing system as outlined by Leutert et al. (2019) without deviation (31). Peptides were digested using sequencing grade trypsin (V5113; Promega) diluted in 50 mM Triethylammonium bicarbonate buffer pH 8.5 (T7408; Sigma). Following digestion, samples were acidified with TFA (A117, Fisher) to a final concentration of 0.5 % (v/v). Peptides were desalted as previously described (32) using an OT-2 liquid handling robot (Opentrons Labworks Inc.) mounted with Omix C18 pipette tips (A5700310K; Agilent). Desalted peptides were dried and stored at −80 °C prior to re-suspension in 3.0 % (v/v) ACN / 0.1 % (v/v) FA for MS injection.

### LC-MS/MS analysis

Peptides were analyzed on a Fusion Lumos Orbitrap mass spectrometer (Thermo Scientific) in data independent acquisition (DIA) mode. A total of 2 μg of re-suspended peptide was injected per replicate using Easy-nLC 1200 system (LC140; Thermo Scientific) and an Acclaim PepMap 100 C18 trap column (Cat# 164750; Thermo Scientific) followed by a 15 cm Easy-Spray PepMap C18 analytical column (ES906; Thermo Scientific) warmed to 50°C. Peptides were eluted at 1.2 μL/min using a segmented solvent B gradient of 0.1 % (v/v) FA in 80 % (v/v) ACN (A998, Fisher) from 4 to 41 % solvent B (0 – 21 min). The FAIMSpro was used with a fixed gas flow of 3.5 L/min with CV settings as described in the manuscript. A positive ion spray voltage of 2.3 kV was used with an ion transfer tube temperature of 300°C and an RF lens setting of 40 %.

### Data Independent Acquisition (DIA)

Direct DIA (dDIA) acquisition was performed as previously described (5). Full scan MS^1^ spectra (350–1400 *m/z*) were acquired with a resolution of 120□000 at 200 *m/z* with a normalized AGC target of 100 % and IT set to automatic. Fragment spectra were acquired at resolution of 30 000 across twenty-eight 38.5 *m/z* windows overlapping by 1 *m/z* using a dynamic maximum injection time and an AGC target value of 2000 %, with a minimum number of desired points across each peak set to 6. HCD fragmented was performed using a fixed 27 % fragmentation energy. BoxCarDIA acquisition was also performed as described (5) using the DIA acquisition outlined above. MS^1^ spectra were acquired using two multiplexed targeted SIM scans of 10 BoxCar windows each. Full scan MS^1^ spectra (350– 1400 *m*/*z*) were acquired with a resolution of 120 000 at 200 *m/z* and normalized AGC targets of 100 % per BoxCar isolation window. Isolation windows used were determined as previously described (5). Fragment spectra were acquired according to the settings described above for dDIA acquisition.

### Data analysis

Downstream data analysis of all dDIA and BoxCarDIA acquisitions was performed on Spectronaut ver. 17 (Biognosys AG) with default settings. Significantly changing differentially abundant proteins (DAPs) were determined and corrected for multiple comparisons (Bonferroni-corrected *p-value* < 0.05; *q-value;* Supplemental Data 1). Gene ontology (GO) analyses were undertaken using The Ontologizer (http://ontologizer.de/; 33) to perform a parent-child enrichment analysis (*p-value* < 0.01). Significantly changing proteins were used as the foreground (Log2FC > 0.58; *q-value* < 0.05) and all quantified proteins as the background. K-means clustering was performed using the *dtwclust* and R ver. 4.2.2 (https://www.r-project.org/) using partitional-type clustering, a ‘dtw’ distance measure, and PAM centroiding. All quantified shoot and root proteins (Supplemental Data 2 and 3) were separated by treatment and imported into Perseus ver. 1.5.16.0 (34). Proteins quantified across all replicates and time-points and possessing a significant change in abundance (BH-corrected ANOVA p-value < 0.05) were selected for z-score conversion and analysis by *dtwclust*. Visualizations were produced using *ggplots2*. Association network analyses were performed using Cytoscape ver. 3.9.0 (https://cytoscape.org/) and the StringDB (35) plugin. Data were supplemented with subcellular localization from SUBA5 (http://suba.live/; 36). Additional plots were produced using GraphPad Prism ver 9 (https://www.graphpad.com/).

### Experimental design and statistical rational

Short gradient BoxCarDIA and dDIA acquisitions were conducted with and without the use of FAIMSpro using the same *Arabidopsis* seedling digest which was generated, aliqoted, dried and frozen at −80 °C prior to use. Characterization of multiple triple-CV settings covering diverse CV ranges were then performed (n = 5; technical replicates) to elucidate an ‘optimal’ setting. *Arabidopsis* roots and shoots subjected to salt and osmotic stress were then analyzed across six time-points beginning at Zeitgeber time 3 (ZT3; 3 h after the onset of light) generating a total of 144 samples (n = 4; biological replicates). We selected 0 h, 1 h, 3 h, 6 h, 12 h, and 24 h time-points post-stress exposure for sampling. Statistical analyses of all proteomic data involved the use of Bonferroni adjusted Student’s t-test to identify significantly changing protein groups (*q-value*) in the shoot and roots samples unless otherwise stated. Bioinformatic analyses consisted of gene ontology enrichment, k-clustering and association network analyses. Figure legends provide further clarification of all statistical tests and corresponding criteria for significance in all analyses.

### Data Availability

Raw data have been deposited to the ProteomeExchange Consortium (http://proteomecentral.proteomexchange.org) via the <u>PR</u>oteomics <u>IDE</u>ntification Database (PRIDE; https://www.ebi.ac.uk/pride/) partner repository with the data set identifiers PXD038989 (Fast-flow DIA dataset) and PXD039004 (Salt/Osmotic Stress dataset).

## Results

### Data Independent Acquisition (DIA) with 3-CV FAIMSpro

To test the capabilities of FAIMSpro in conjunction with a short 21-minute fast-flow gradient, we utilized two established single-shot DIA acquisition workflows; dDIA (37) and BoxCarDIA (5) to analyze *Arabidopsis* seedling extract. Here, we head-to-head compared multiple 3-CV FAIMSpro settings using each acquisition approach (Figure 1). We selected four 3-CV ranges on the basis of previous literature (38; Figure 1) and found that a FAIMSpro CV setting of −30 −50 −70 produced the largest number of quantified protein groups. Further, when a −30 −50 −70 FAIMSpro setting was used in conjunction with dDIA, it outperformed BoxCarDIA in the overall number of quantified protein groups (Figure 2*A*; Supplemental Data 1). Here, a total of 5540 and 5162 protein groups were quantified by dDIA and BoxCarDIA, respectively (Figure 2*A*, Supplemental Data 1). A more contracted 3-CV setting of −40 −55 −75 narrowed the gap between these two acquisition approaches in terms of overall quantified protein groups, with 5445 and 5400 protein groups quantified by dDIA and BoxCarDIA, respectively (Figure 2*A*; Supplemental Data 1).

**Figure 1.**
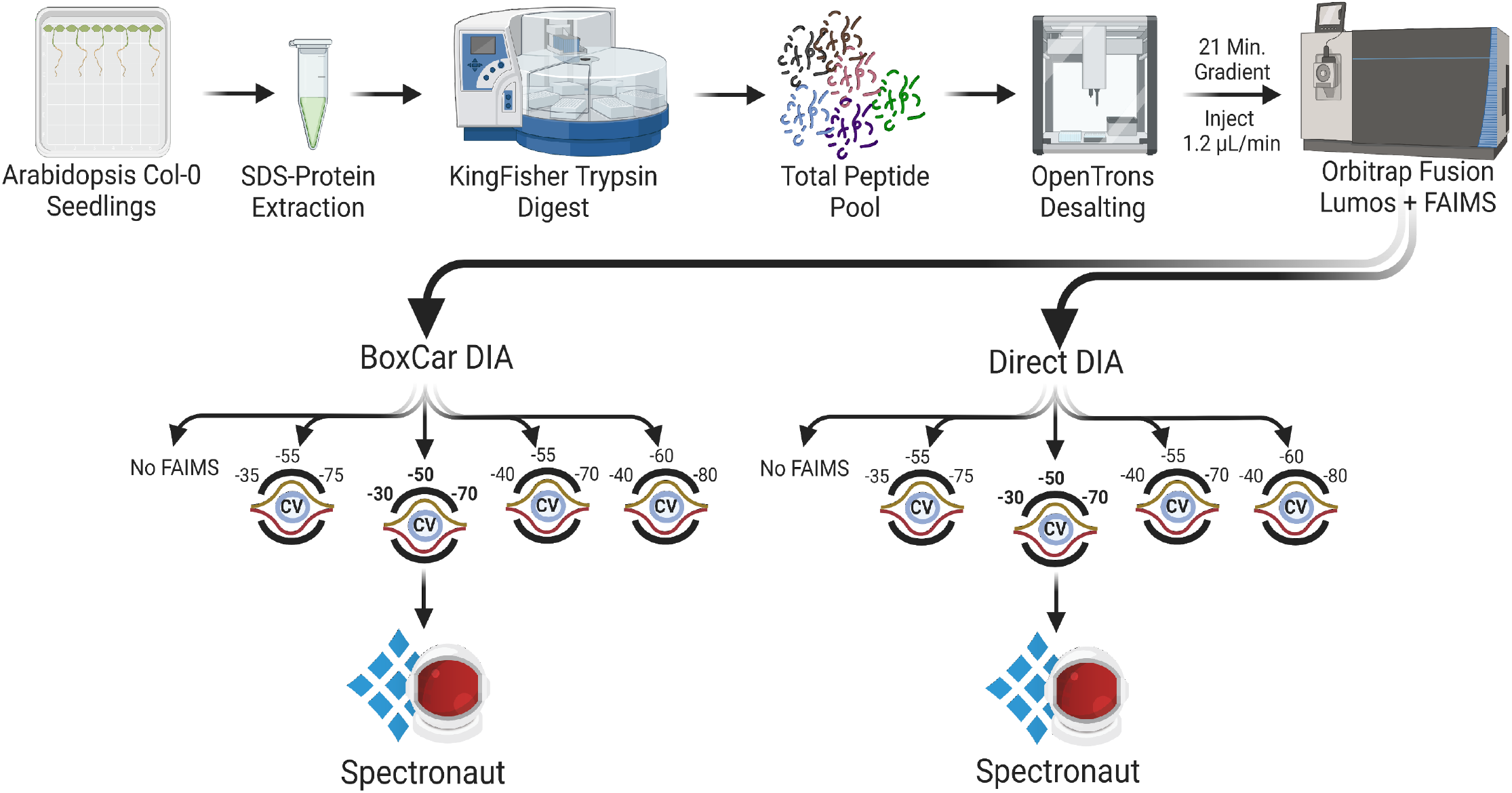
Fast-flow short-gradient DIA acquisition workflow. *Arabidopsis* Col-0 seedlings were extracted and digested as previously described (5), followed by desalting using an OpenTrons OT-2 liquid handling robot. Replicate samples were resolved over a 21-minute micro-flow gradient and subjected to BoxCarDIA or DirectDIA acquisition on an Orbitrap Fusion LUMOS with or without a FAIMSpro device (n = 5). Multiple Multi-CV FAIMS acquisitions were performed, with all acquired DIA data processed using Spectronaut v17 (Biognosys, AG).

**Figure 2.**
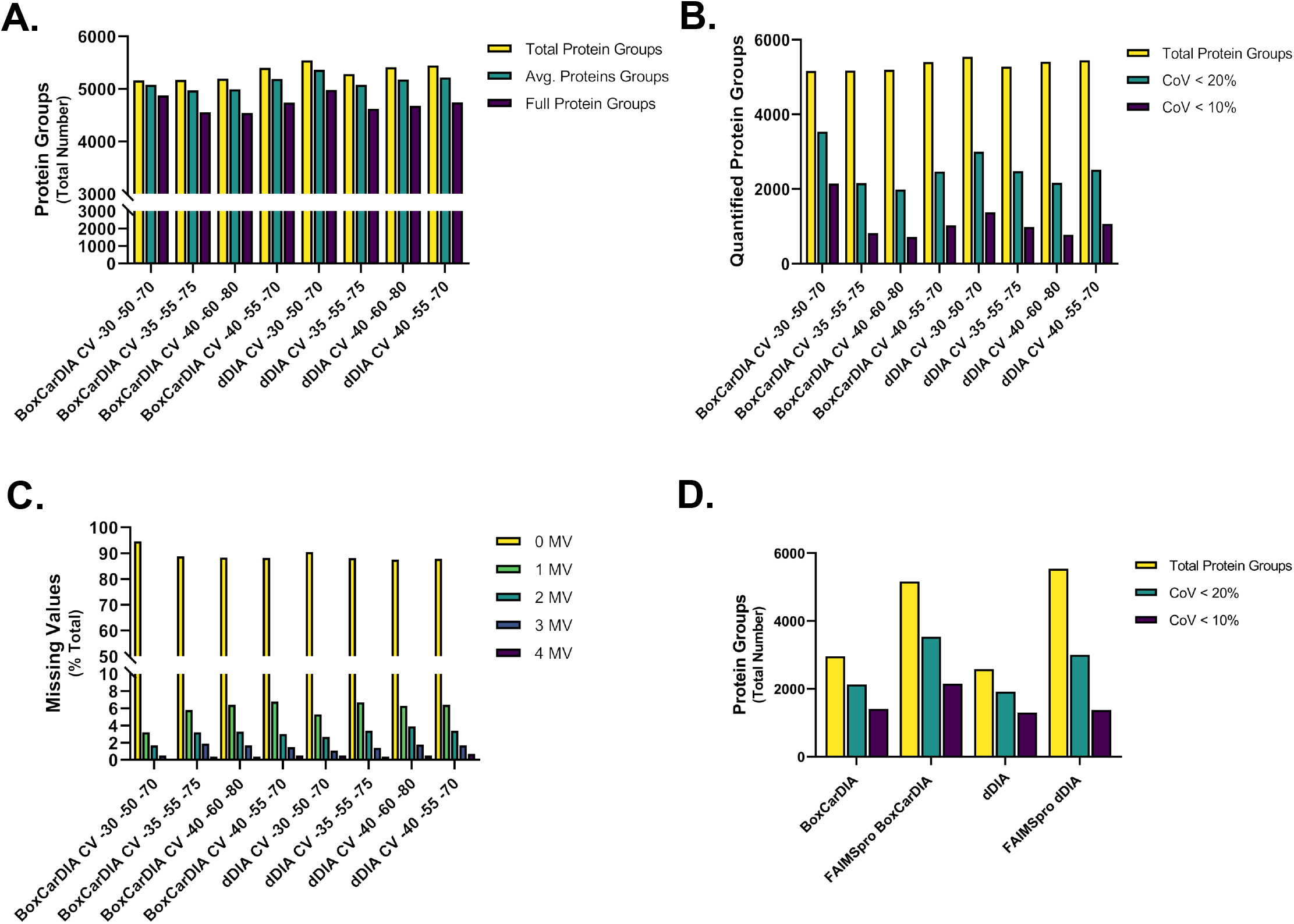
Short-gradient multi-CV FAIMSpro BoxCarDIA acquisition. Injections consisting of 2 μg *Arabidopsis* seedling peptide each were performed for each 21-minute FAIMSpro DIA acquisition (n=5). *A*. Total, average and full protein groups acquired by either BoxCarDIA or DirectDIA across five injections using different multi-CV FAIMSpro settings. *B*. Total protein groups quantified, protein groups with a coefficient of variance (CoV) less than 20% and less than 10% for each DIA acquisition method. *C*. Percent missing values (MV) for each DIA acquisition method performed. Missing values range from 0 to 4. *D*. Total protein group quantification and CoV differences between fast flow BoxCarDIA and DirectDIA acquisitions performed with FAIMSpro (CVs −30, −50, −70) and without FAIMSpro.

Next, we assessed all four 3-CV FAIMSpro settings in combination with either dDIA or BoxCarDIA for their quantitative accuracy by analyzing the reproducibility of protein intensities between replicates (n = 5). The 3-CV −30 −50 −70 FAIMSpro setting yielded the best co-efficient of variance (CoV) when using either dDIA or BoxCarDIA acquisition methods, with 58 % and 69 % of all protein groups exhibiting a CoV < 20 %, respectively (Figure *2B;* Supplemental Data 1). This FAIMSpro BoxCarDIA CoV value is comparable to previously obtained single-CV FAIMS dDIA HeLa digest data, where 71 % of ~5100 protein groups obtained over a 21-minute gradient possessed a CoV < 20 % (3). Despite the ability of dDIA to quantify more proteins groups, it is ~11 % less accurate in a 3-CV FAIMSpro configuration when analyzing *Arabidopsis* seedlings relative to BoxCarDIA (Figure 2*B*; Supplemental Data 1).

Missing values (MV) represent a major limitation in MS analyses and continue to be a critical criteria for the development of novel acquisition methods (5). As such, we queried our 3-CV FAIMSpro dDIA and BoxCarDIA acquisitions for MV (Figure 2*C*; Supplemental Data 1). Using a FAIMSpro 3-CV setting of −30, −50, −70 we find that 90.5 % and 94.6 % of the quantified dDIA and BoxCarDIA protein groups to possess no MV (Figure 2*C*; Supplemental Data 1). However, despite quantifying more protein groups, dDIA maintains nearly twice as many proteins groups with 1 – 3 MV compared to BoxCarDIA (9.1% vs. 5.4%), along with five times the number of proteins groups with four MV (Figure 2*C*; Supplemental Data 1). Lastly, using a FAIMSpro 3-CV setting of −30 −50 −70 with either dDIA or BoxCarDIA, we compared the overall improvement in quantified protein groups provided by using FAIMSpro with 3-CVs versus no FAIMSpro (Figure *2D;* Supplemental Data 1). We find that FAIMSpro provides a substantial improvement in overall quantified protein groups ranging from a 42.7 % (BoxCarDIA) to 53.4 % (dDIA) increase (Figure 2*D*; Supplemental Data 1). Overall, our data indicate that FAIMSpro substantially increases the total protein groups quantified by both dDIA and BoxCarDIA. However, despite quantifying 6.8 % (378 total) more protein groups by dDIA, BoxCarDIA demonstrated lower overall CoVs and MVs relative to dDIA when analyzing *Arabidopsis* seedlings.

#### Time-course analysis of Arabidopsis seedlings subjected to salt and osmotic stress

The ability to execute quantitative proteomic time-course experimentation in plants is restrained by a number of factors. These include: 1) The compromise between sensitivity and analysis speed due to the high dynamic range and complexity of plant samples and 2) the lack of accessible MS instrumentation to execute quantitative proteomic experimentation. Our development of a fast-flow, short-gradient 3-CV FAIMSpro BoxCarDIA methodology for the analysis of plant samples represents a means to overcome these obstacles, with our workflow offering the opportunity to analyze 3 – 4x more samples per day at a depth of analysis comparable to a standard 2 h LC-MS/MS workflow. We therefore applied our fast-flow, short-gradient FAIMSpro BoxCarDIA acquisition approach using a −30 −50 −70 3-CV setting to perform quantitative proteomic profiling of *Arabidopsis* seedlings subjected to salt and osmotic stress over a 24 h period (Figure 3). Across our time-course we were able to quantify >6400 shoot (Supplemental Data 2) and >8500 root (Supplemental Data 3) protein groups, equaling 9647 total uniquely quantified protein groups between these two plant organs.

**Figure 3.**
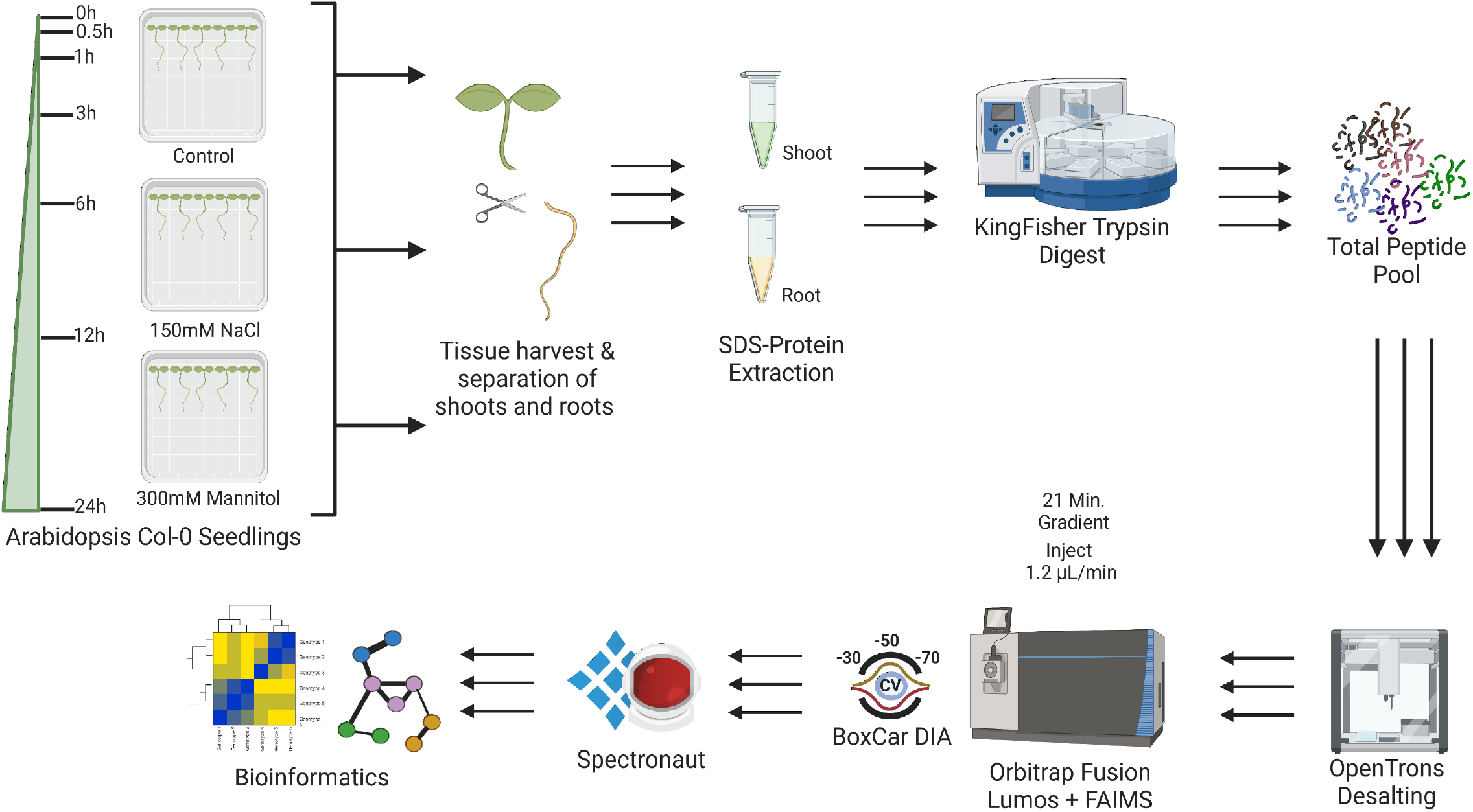
Application of fast-flow multi-CV FAIMSpro BoxCarDIA to detect *Arabidopsis* proteome changes in response to salt and osmotic stress. *Arabidopsis* Col-0 seedlings were grown on 0.5x MS media for 14 days prior to transfer to fresh 0.5x MS plates maintain control, salt (150mM NaCl) or osmotic (300mM Mannitol) conditions for up to 24h. Upon harvesting, shoots and roots were separated prior to extraction, digestion and FAIMSpro BoxCarDIA analysis as outline in Figure 1 (n = 4).

#### Gene Ontology analysis reveals organ specific protein responses

The evaluation of shoots and roots found a large number of Differentially Abundant Proteins (DAPs) (Log2FC >0.58; *q-value* < 0.05) under both salt (150 mM sodium chloride; NaCl) and osmotic (300 mM mannitol; Man) stress conditions. Interestingly, we found that across all time-points and within each organ, post-stress initiation, there are relatively few overlapping DAPs between salt and osmotic stress responses. Such results suggest that although salt and mannitol are commonly considered as related abiotic stresses that lead to similar biological process perturbations, at the proteome-level there are more stress-specific cellular responses than shared responses (Figure 4*A*; Supplemental Data 4).

**Figure 4.**
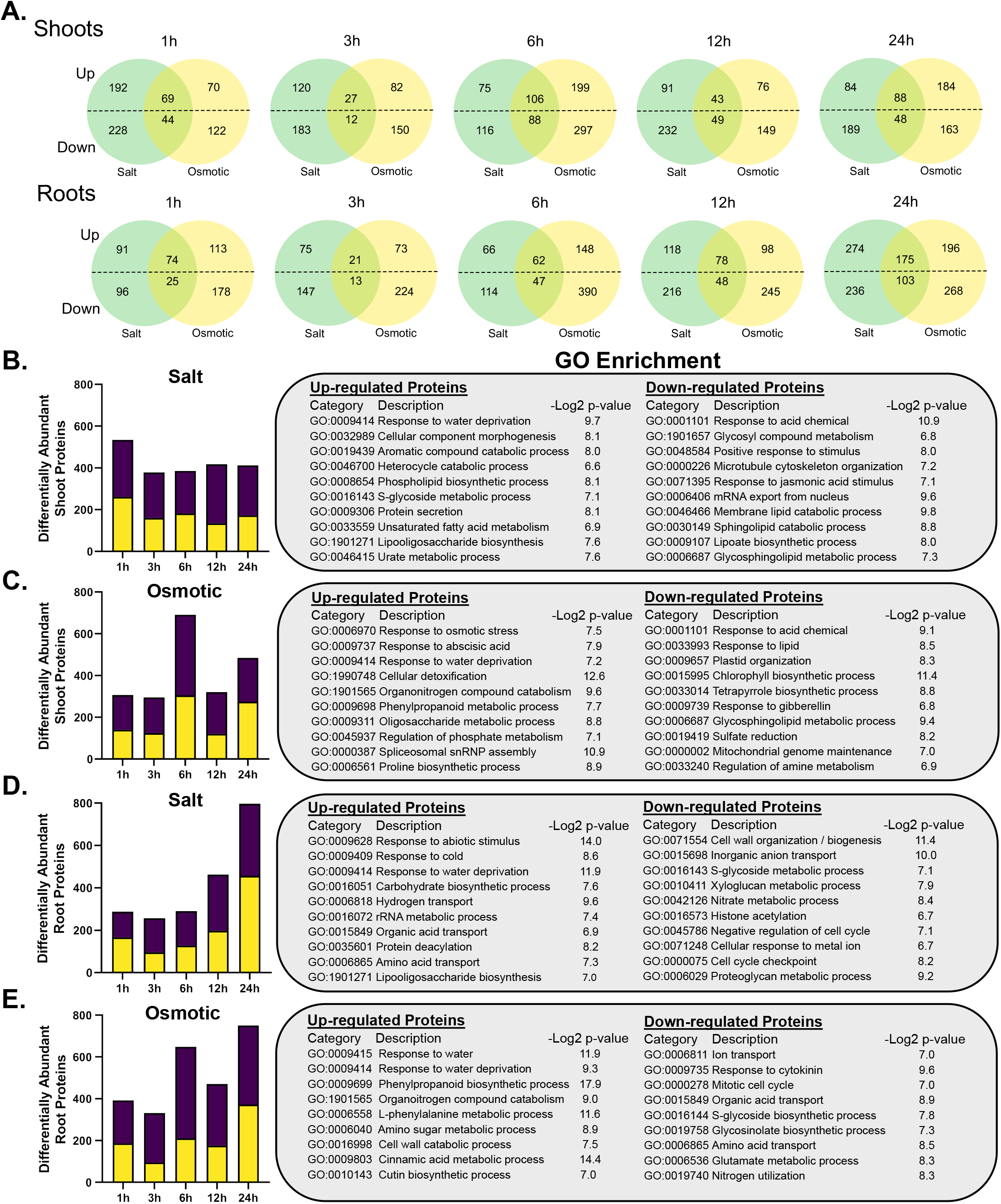
Time-course proteome analysis of salt and osmotic stress responses in *Arabidopsis* shoots and roots. *Arabidopsis* Col-0 seedlings were grown on 0.5x MS media for 14 d prior to transfer to fresh 0.5x MS plates maintain control, salt (150mM NaCl) or osmotic (300mM Mannitol) conditions for up to 24 h. *A*. Venn diagram analysis of the significantly changing up- and down-regulated proteins in either shoots or roots at each time-point. Proteins with overlapping responses for treatment at each time-point is shown (Log2FC > 0.58; *q-value* < 0.05). *B-C*. shoot and *D-E*. root differential proteome responses to salt and osmotic stress, respectively. The total, significantly changing up (yellow) and down (purple) regulated proteins for each time-point (n = 4, Log2FC > 0.58; *q-value* < 0.05). Corresponding enriched gene ontology (GO) categories for up and down-regulated proteins are shown for each treatment across the time-course (*p-value* < 0.01; −Log2 *p-value* depicted).

Given this, we next examined stress-specific up- and down-regulated proteins in both shoots and roots for the enrichment of biological processes. Here, we find a number of DAPs to be related to osmotic / salt stress-associated gene ontology (GO) categories (Figure 4 *B-E;* Supplemental Data 5), such as *Response to Water Deprivation* (GO:0009414), which was enriched amongst up-regulated DAPs in both osmotic and salt treated shoot samples (Figure 4 *B-C*). Likewise, we find *Cellular Response to Metal Ion* (GO:0015698) and *Ion Transport* (GO:0006811) enriched amongst down-regulated DAPs in roots following exposure to salt and mannitol, respectively, which is consistent with both stressors disrupting ion homeostasis in plant cells (Figure 4 *D-E*; 39). In addition, GO categories *Sphingolipid Catabolic Process* (GO:0030149) and *Lipid-Related Processes* (GO:0006687, GO:000149 and GO:0006687) were enriched in up- and down-regulated DAPs, respectively, in both salt and osmotic stress treated shoot samples, indicating dynamic compositional changes occurring in the shoot cell membrane over 24 h of stress. Consistent with canonical root responses to salt and osmotic stress, we also find cell wall related GO categories enriched amongst down-regulated DAPs in root samples (Figure 4 *D-E*; 39, 40). These include: *Cell Wall Organization / Biogenesis* (GO:0071554) and *Cell Wall Catabolic Process* (GO:0016998). Interestingly, we find that phytohormone responses differ between the shoots and roots with *Response to Abscisic Acid* (GO:0009737) and *Response to Gibberellin* (GO:0009739) enriched amongst the up- and down-regulated DAPs in shoots subjected to osmotic stress (Figure 4 *B-C*), while *Response to Cytokinin* (GO:0009735) is enriched amongst down-regulated DAPs in root tissues under osmotic stress. Together, these results indicate that under abiotic stress conditions, phytohormone pathway responses differ at the protein-level in shoots and roots (Figure 4 *B-E*).

#### Time-resolved systems-level changes in the shoot and root proteome

To highlight our study’s ability to elucidate time-resolved protein-level responses upon exposure to salt or osmotic stress, we specifically focused on two conditions; mannitol-exposed roots (Figure 5; Supplemental Data 6,8) and salt-treated shoots (Figure 6; Supplemental Data 7,9), each of which contained the largest number of up-regulated DAPs (BH-corrected *p-value* < 0.05). First, we conducted a K-mean clustering analysis of all DAPs from mannitol treated roots, which resolved 4 x DAP clusters. Cluster 1 DAPs peak 1 h after stress onset, suggesting they have a role in establishing osmotic stress signaling (Figure 5 *A-B*). This is highlighted by enrichment of *Molybdate Ion Transport* (GO:0015689), as molybdate represents a key co-factor for ABA biosynthetic enzyme ABA DEFICIENT 3 (ABA3), which is responsible for ABA production and anthocyanin accumulation (42, 43). Cluster 2 DAPs are down-regulated immediately after stress initiation, indicating that their response may be attenuated upon stress exposure (Figure 5*B*). Here, we find enrichment of *Cell Growth* (GO:0016049), indicating that plants are reducing growth in order to allocate resources to stress response (44). Rather than peaking at multiple time-points, as in seen Cluster 3, Cluster 4 exhibits a gradual increase in abundance over 24 h of exposure to osmotic stress, suggesting that these DAPs are required to mitigate the on-going osmotic stress and that these DAPs likely have direct roles in protecting plants from the damaging effects of extended stress exposure.

**Figure 5.**
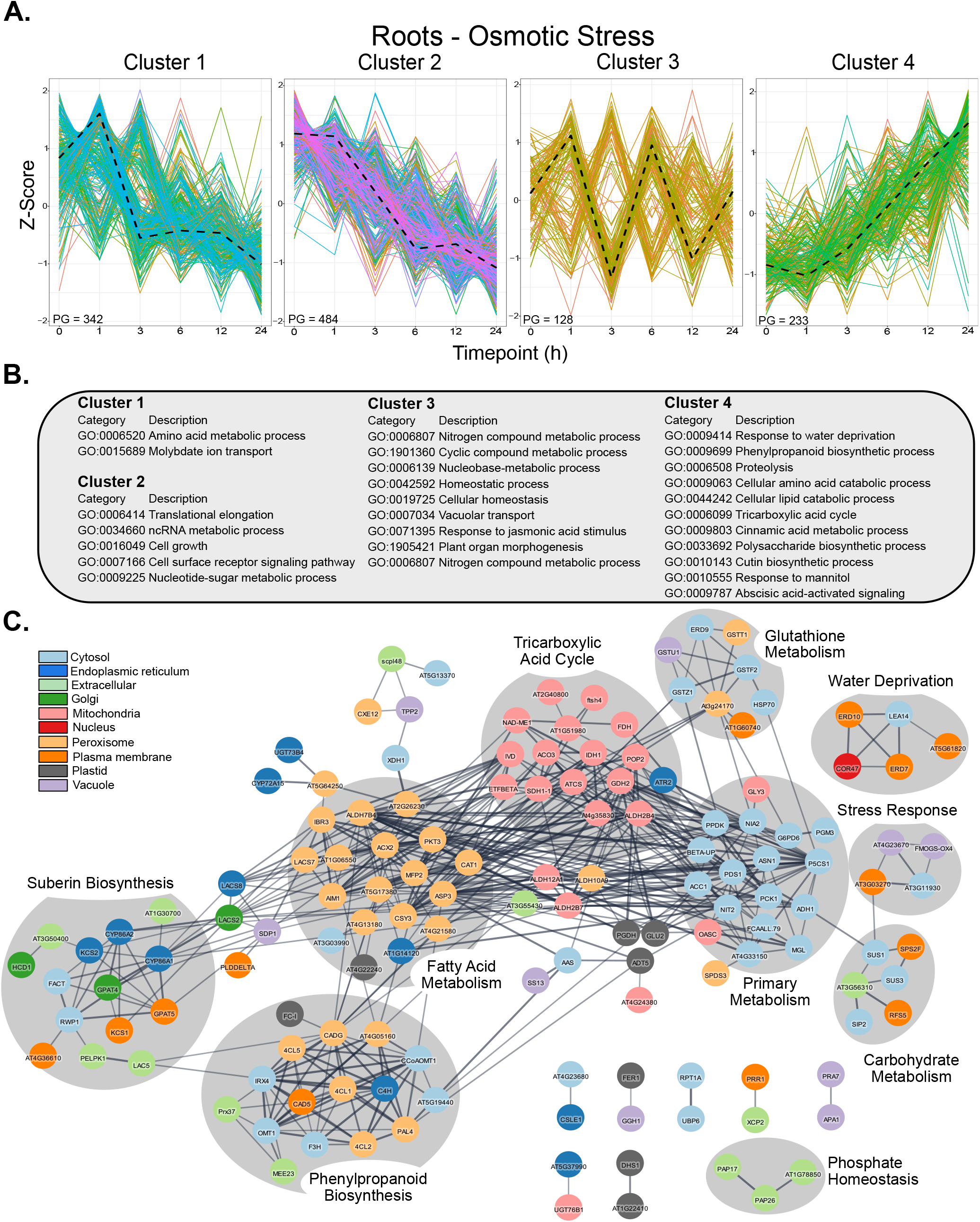
Proteome-level changes in osmotic stressed roots over 24h. *A*. K-cluster analysis of significantly changing proteins induced by osmotic stress in *Arabidopsis* roots over a 24h time-course (z-score; Benjamini-Hochberg (BH)-corrected ANOVA *p-value* < 0.05). *B*. Select enriched GO biological process categories of cluster 1 through 4 (*p-value* < 0.01; Supplemental Data 8). *C*. STRING-DB association network of cluster 4 proteins with edge scores > 0.6. Nodes with edges below this threshold were removed. Node colors represent subcellular localization (SUBAcon); see legend. Highlighted node clusters were categorized with assistance from the STRING-DB functional annotation.

Cluster 4 is the most likely to contain proteins of interest for the targeted breeding of improved drought tolerance in crops, thus we further contextualized these proteins and their potential cellular relationships through a STRING-DB association network analysis (Figure 5*C*). Here, we find a number of stress-related protein clusters, including proteins from water deprivation responses (e.g., EARLY RESPONSIVE TO DEHYDRATION 10 (ERD10, AT1G20450)) (45). In addition, we also found several aspects of metabolism to be altered. For example, we find a protein cluster related to suberin biosynthesis (Figure 5*C*), which is consistent with suberin being reported to prevent root water loss (46, 47). We also find phenylpropanoid biosynthetic proteins to be a large portion of Cluster 4 (Figure 5*C*), with their accumulation having been related to abiotic stress resistance in apple (48) and wheat (49). Tricarboxylic Acid (TCA) cycle proteins are also abundant in Cluster 4 and have been related to root growth via secondary cell wall generation (50). In addition, we also see large clusters of carbohydrate and glutathione metabolism proteins, which have intimate relationships with abiotic stress resistance in *Arabidopsis* (51–54). Lastly, we examined the subcellular localization distribution of Cluster 4 DAPs to resolve which cellular compartments are most affected in roots. We find that Cluster 4 DAPs are predominantly cytosolic (e.g. primary metabolism), mitochondrial (e.g. TCA cycle) and peroxisomal (e.g. fatty acid metabolism), with relatively minimal protein abundance changes observed in proteins targeted to the nucleus and plastid.

Next, we examined the response of shoots to high salinity conditions (Figure 6). We again resolved four DAP clusters. Cluster 1 represents an early responding cluster, which contains DAPs involved in *Carbohydrate Metabolic Processes* (GO:0005975) and *Response to Oxidative Stress* (GO:0006979). Cluster 2 contains DAPs that peak at 3 h and 12 h, demonstrating a prolonged response. Cluster 3 contains DAPs generally down-regulated in response to salinity stress, while Cluster 4 included intermediate responding DAPs that peak at 3 h post-stress initiation (Figure 6*A*).

**Figure 6.**
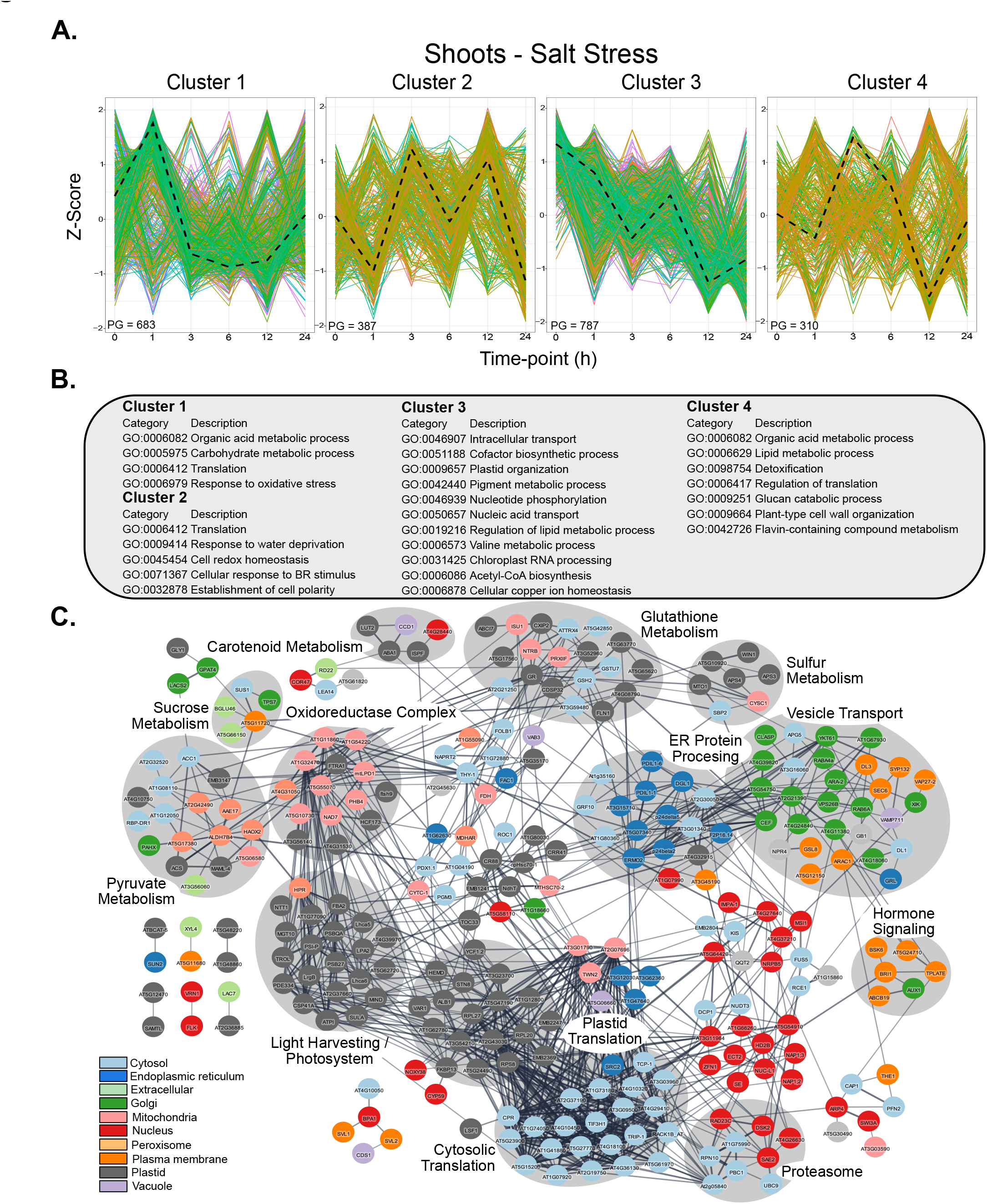
Proteome-level changes in salt stressed shoots over 24h. A. K-cluster analysis of significantly changing proteins induced by salt stress in *Arabidopsis* shoots over a 24h time-course (z-score; Benjamini-Hochberg (BH)-corrected ANOVA *p-value* < 0.05). *B*. Select enriched GO biological process categories of cluster 1 through 4 (*p-value* < 0.01; Supplemental Data 9). *C*. STRING-DB association network of cluster 2 proteins with edge scores > 0.6. Nodes with edges below this threshold were removed. Node colors represent subcellular localization (SUBAcon); see legend. Highlighted node clusters were categorized with assistance from the STRING-DB functional annotation.

With Cluster 2 (Figure 6*A*), containing the largest number of up-regulated proteins over multiple time-points, we performed GO and STRING-DB analyses to contextualize the protein constituents and their involvement in different biological processes. Here, we find a number of enriched GO categories, such as *Protein Translation* (GO:0006412), *Response to Water Deprivation* (GO:0009414) and *Cell Redox Homeostasis* (GO:0045454), which we see reflected as discernable protein clusters in our STRING-DB association network analysis. Similar to mannitol-exposed root cells, we also find a large protein group related to glutathione metabolism, indicating that increases in glutathione metabolism is a common abiotic response mechanism at the protein-level in both shoots and roots over the first 24 h of stress. We also see other facets of metabolism emerge as protein clusters, including sucrose (55, 56), carotenoid (57, 58), pyruvate (59, 60) and sulfur metabolism (61, 62), each of which have been implicated in plant abiotic stress resistance to varying degrees. Unlike the most comparable cluster from mannitol treated roots (Cluster 4), we see DAPs localized to a wide diversity of subcellular localizations, highlighted by large clusters of cytosolic- and plastid translation-related DAPs. Interestingly, unlike mannitol stressed roots, we see relatively few peroxisome-localized DAPs, but instead find a proportional increase in nuclear localized DAPs.

## Discussion

The need to effectively perform time-course or time-series proteomic experiments is critical to developing a wholistic understanding of the dynamically changing plant cell landscape. However, this comes with substantial acquisition time overheads, which has limited proteomic advancements in the molecular plant sciences when compared to RNA sequencing technologies. As such, we developed a method utilizing multi-CV FAIMSpro and a short 21-minute gradient in order to increase the speed of acquisition without limiting depth of analysis. When coupled with fast-flow liquid chromatography gradients, single-CV FAIMSpro DIA was able to increase the number of samples analyzed per day at a comparable depth of analysis to longer gradients (3, 63). However, the use of multiple CVs has surpassed the single-CV approach due to its ability to select a wider range of precursors for MS analysis in a single injection. Our comparison of dDIA (37) and BoxCarDIA (5) acquisition approaches with and without FAIMS, found that multi-CV FAIMSpro BoxCarDIA methods were superior in CoV, while also possessing fewer MVs, confirming previous reports using 2 h gradients (5) and further indicating the benefits of MS^1^ segmentation when performing orbitrap-based DIA (5, 7). Owing to the nature of plant samples we also found a substantial increase in the overall number of quantified proteins when using FAIMSpro, empirically demonstrating its utility when analyzing plant samples by orbitrap-based DIA. This supports DDA-based multi-CV FAIMSpro acquisitions, which have demonstrated increases in both quantified PTM events (64, 65) and protein abundance (38) changes.

### Temporal analysis of salt and mannitol induced proteome changes in Arabidopsis reveals unique response trajectories

Multiple time-course experiments have described transcriptomic changes triggered by osmotic or salt-stress in *Arabidopsis* (66, 67), with none endeavoring to extensively quantify temporal protein-level changes. Proteins represent the effectors of the plant cell, rendering the lack of time-resolved quantitative proteomic data relating to their salt and osmotic stress responses a critical gap in our understanding. Interestingly, plants are proposed to have three stages of abiotic stress response after stress onset, including: a stop phase, quiescent phase and a recovery phase (68, 69), during which, plants modify their growth rate to allocate resources for stress tolerance (70). For example, 5 d-old *Arabidopsis* seedlings exposed to salt stress revealed that 4 to 5 h after treatment, growth rates rapidly slowed down (stop phase), remaining steady for an additional 4 h (quiescent phase) before recovering growth 13 to 24 h after stress onset (69). During this response reprogramming, plants undergo extensive gene expression changes to temporally regulate and attenuate the plant cell response to abiotic stress. However, what happens temporally at the protein-level and across different organs remains unknown.

Therefore, we applied salt and mannitol as abiotic stressors, which are two stimuli commonly considered to have overlapping responses, such as the induction of ABA and reduction of primary root elongation. Surprisingly, at the protein-level we find more non-overlapping DAPs induced by salt and mannitol than overlapping DAPs (Figure 4*A*). Apart from the common negative impacts of hyperosmotic stress triggered by salt and mannitol treatments, a salt treatment imposes additional ionic stress (71), with plants compartmentalizing Na^+^ into the vacuoles to maintain cellular ion homeostasis and turgor pressure (72). This is highlighted in both organs where salt stress induces more overall DAPs than osmotic stress. However, at the organ-level we find salt stress induces more down-regulated DAPs than osmotic stress in shoots, while the opposite is observed in roots. In only one instance (up-regulated DAPs in shoots at 6 h) do we find more overlapping DAPs than uniquely up-regulated salt induced DAPs. Why *Arabidopsis* exhibits these differential organ responses likely stems from the different environmental challenges faced by these organs under abiotic stress (e.g. the presence of light), however, fully resolving this is beyond the scope of the current study.

### Lipid-related biological processes dominate enriched gene ontologies across organs

Across our time-course experiment, we find the enrichment of multiple GO categories relating to lipid metabolism in shoots under either salt or osmotic stress, including an up-regulation of *Phosphoplipid Biosynthetic Processes* (GO:0008654) and the down-regulation of *Sphingolipid Catabolism* (GO: 0030149) and *Glycosphingolipid Metabolic Processes* (GO:0006687). Phospholipids are well-characterized constituents of the bi-layer membrane (73). Studies have shown that phospholipids, such as phosphatidic acid (PA), are important for fine-tuning stress signaling pathways through allosteric protein binding (73, 74). Abiotic stress is also shown to widely affect phospholipid content in *Arabidopsis* (75), along with tobacco (*Nicotiana tabacum;*75), rice (Oryza sativa; 76) and the common ice plant (Mesembryanthemum crystallinum; 77). Sphingolipids however, have only recently been implicated in abiotic stress response through proteins like ACCELERATED CELL DEATH 5 and 11 (ACD5; AT5G51290 and ACD11; AT2G34690)(79, 80), with ACD11 protein levels up-regulated in seedlings exposed to 150 mM salt over 24 h and constitutive over-expression of ACD11 improving plant salt and drought resistance (79). While we find ACD11 proteins down-regulated in roots after 3 to 6 h of mannitol treatment, we do not see changes in either ACD5 or 11 in the shoots. Further, we find the down-regulation of sphingolipid related proteins DIHYDROSPHINGOSINE PHOSPHATE LYASE 1 (DPL1; AT1G27980), GLCCER SYNTHASE (GCS; AT2G19880), and two GBA2-type family beta-glucosidases (AT3G24180 & AT5G49900) in shoots, which are related to *Sphingolipid Catabolism* (GO: 0030149) and/or *Glycosphingolipid Metabolic Processes* (GO:0006687). This indicates there may be unique protein-level organ-specific roles for sphingolipid-related metabolism in response to salt and osmotic stress.

Enrichment of *Lipid Catabolic Processes* (GO:0044242) was also found in Cluster 4 of osmotic stressed roots (Figure 5 *A-B*), suggesting that lipid breakdown increases over time in roots under osmotic stress (Figure 5*B*). This is supported by previous phospholipase research, which found that phospholipid hydrolysis can promote stress responses in *Arabidopsis* (81, 82). Further, our Cluster 4 association network resolved a large network of peroxisome-targeted fatty acid metabolism proteins, which are intimately linked with lipid homeostasis (Figure 5*C*). Interestingly, almost none of these proteins have been investigated in an abiotic stress context. Collectively, these lipid metabolism data highlight that broad-ranging organ-specific differences exist at the proteome-level in response to abiotic stress.

### Time-resolved proteome changes induced by salt and mannitol stress reveals multiple links between metabolism and abiotic stress response

To mitigate abiotic stress, plants modify gene expression, protein abundance, and PTMs as well as the accumulation of metabolites (83). Unlike other studies, which have explored end-point proteome-level analyses (21, 84, 85), our new fast-flow multi-CV FAIMSpro BoxCarDIA workflow enabled us to derive a time-resolved understanding of the abiotic stress proteome and uncover several organ-specific links between plant metabolism and abiotic stress response. We highlight this by examining the clusters of DAPs in roots and shoots that exhibit an increase in abundance over time in response to abiotic stress. Here, we find phenylpropanoid metabolism (roots), suberin biosynthesis (roots), vesicle transport (shoots), carotenoid biosynthesis (shoots) and glutathione metabolism (roots and shoots) as critical factors, in addition to primary metabolic changes that respond to either osmotic stress in roots or salt stress in shoots.

We specifically find suberin metabolism enriched in root tissues following exposure to osmotic stress. Suberin, being a hydrophobic polymer, is deposited with lignin to the endodermis barrier, therefore affecting water and nutrients transported into roots from soil (86, 87). In addition, when responding to high salinity, plants can accumulate suberin in order to prevent water loss and Na^+^ absorption (88–90). In our analysis, we see a number of proteins related to suberin biosynthesis enriched in Cluster 4 mannitol-treated root samples (Figure 5*C*). In particular, 3-ketoacyl CoA synthase 2 (KCS2; AT1G04220), REDUCED LEVELS OF WALL-BOUND PHENOLICS 1 (RWP1; AT5G41040), and FATTY ALCOHOL:CAFFEOYL-COA CAFFEOYL TRANSFERASE (FACT; AT5G63560), of which KCS2 and FACT have been previously related to osmotic and salt stress resistance, respectively (91, 92).

Upstream of suberin production is phenylpropanoid biosynthesis, which we also find to be an abundant cluster of DAPs (Figure 5*C*). Phenylpropanoids can be converted to diverse aromatic metabolites, such as lignin, suberin, and as well as flavonoids/anthocyanins, therefore impacting virtually all aspects of plant development and fitness (93–95). In terms of abiotic stress responses, phenylpropanoid biosynthesis is linked to cell wall thickening through lignin accumulation (96) and enhanced anti-oxidative activity through the production of phenolics (97). However, a direct connection between phenylpropanoid metabolism and osmotic stress responses remains elusive, at least at the protein-level. Here we see that osmotic stress up-regulates a set of phenylpropanoid biosynthetic proteins, including: PHENYLALANINE AMMONIA-LYASE 4 (PAL4; AT3G10340), 4-COUMARATE-CoA LIGASE (4CL1; AT1G51680, 4CL2; AT3G21240 & 4CL5; AT3G21230), and CAFFEOYL-CoA 3-O-METHYLTRANSFERASE (CCoAOMT1; AT4G34050) (98). Among these, expression of CCoAOMT1 was found to be induced by both mannitol and salt treatments (99, 100), while our data shows a similar increase in CCoAOMT1 protein levels in response to osmotic stress. Additionally, PAL proteins are responsible for catalyzing the first step of phenylpropanoid pathway and responding to environmental stress (101), with PAL1 and 2 proposed to be the principal functional isoforms in plants (98, 102). However, we found PAL4 (AT3G10340) up-regulated in response to osmotic stress over 24 h, which is supported by the observation that PAL4 is transcriptionally induced by other abiotic stress stimuli, such as cold treatments (103). This implies that the various PAL proteins may all be functional enzymes, but activated under varying environmental conditions. Beyond this, we resolved a wide range of uncharacterized osmotic stress-involved phenylpropanoid synthesis proteins (Figure 5*C*), providing a direct intersection between phenylpropanoid metabolism and osmotic stress resistance.

In *Arabidopsis* shoots, we uncovered a network of proteins related to vesicular trafficking following exposure to salt stress (Figure 6*C*). Vesicle transport is generally achieved by two preliminary pathways; secretory and endocytic (104, 105). Newly synthesized proteins coupled with exocytic vesicles from endoplasmic reticulum are secreted to the Golgi apparatus and the transGolgi network before being released to various destinations, such as plasma membrane (PM) and vacuole (106, 107). Endocytosis directs internalization of PM proteins from PM into cytoplasm (108, 109). As fundamental processes in plants and other eukaryotic organisms (110), endocytosis and exocytosis have been linked to abiotic stress. For example, components of ABA signaling, ranging from perception to transduction, were found to be controlled by the endocytic trafficking pathway (111, 112). However, knowledge of the proteins responsible for vesicular transport under abiotic stress remain limited (113–116). A previous study showed that over-expression of VESICLE-ASSOCIATED MEMBRANE PROTEIN 711 (VAMP711; AT4G32150) boosts drought stress by dampening PM H^+^-ATPase activity (116), consistent with our results that VAMP711 protein levels are up-regulated in response to salt exposure (Figure 5*C*). In addition, DYNAMIN-LIKE PROTEIN 1 and 3 (DL1; AT5G42080 & DL3; AT1G59610) influence biotic stress responses by affecting endocytosis of the immune receptor FLAGELLIN SENSING2 (FLS2; AT5G46330), although roles of DL1 and DL3 in abiotic stress remains unknown, these two proteins are up-regulated in our salt-treated roots analysis. Further, research exploring the roles of these DL proteins, amongst the many other that we find to be up-regulated, represent new and exciting avenues for future research into mitigating drought-related stresses.

In salt-treated shoot tissues, we also found carotenoid metabolism to be up-regulated (Figure 6 *A,C*). Carotenoids are potent antioxidants, also functioning as an oxidative stress signal in plants (117, 118). In addition, carotenoid biosynthetic genes facilitate ABA accumulation during salt stress (58). In particular, we find CAROTENOID CLEAVAGE DIOXYGENSE 1 (CCD1; AT3G63520), which has been reported to be induced by abiotic stressors in poplar (119) and pepper (120) as well as ABA DEFICIENT 1 (ABA1; AT5G67030), which encodes the enzyme catalyzing the first committed step of ABA biosynthesis (121). Interestingly, beyond CCD1, we see up-regulation of other xanthophyll biosynthetic enzymes such as LUTEIN DEFICIENT 2 (LUT2; AT5G57030) and 2C-METHYL-D-ERYTHRITOL 2,4-CYCLODIPHOSPHATE SYNTHASE (ISPF; AT1G63970), which remain uncharacterized in relation to abiotic stress response.

Lastly, we find a glutathione metabolism DAP networks in both salt-stressed shoots and osmotic-stressed roots (Figure 5*C* and 6*C*). Due to its roles as a thiol antioxidant as well as a reactive oxygen species (ROS) scavenger, accumulating evidence suggests that increased exogenous and reduced endogenous levels of glutathione could boost abiotic stress resistance in *Arabidopsis* (52, 122). Our analyses show that both salt-treated shoots and mannitol-treated roots exhibit changes in glutathione metabolism, which aligns with an increased need to scavenge ROS induced by either abiotic stress (123). Interestingly, we find different DAPs related to glutathione metabolism in salt-exposed shoots and mannitol-treated roots, suggesting that plants might produce, accumulate, and utilize glutathione in an organ specific manner through distinct protein isozymes or regulatory mechanisms. For example, GLUTATHIONE S-

TRANSFERASE TAU CLASS 1 (GSTU1; AT2G29490) and GLUTATHIONE S-TRANSFERASE PHI CLASS 2 (GSTF2; AT4G02520) were found up-regulated in osmotic stressed roots, while GSTU7 (AT2G29420) was found up-regulated in shoots, suggestive of specificity within the GST protein class. So far, only a handful of glutathione S-transferases (GSTs) have been characterized in *Arabidopsis* (122, 124, 125). Therefore, proteins, such as GSTU1 and GSTF2, could offer further insight into how GSTs fine-tune abiotic stress adaptions. Our protein-level results demonstrating that glutathione metabolism enzyme abundance peaks at 3 h, and then recovers to pre-stress levels by 24 h of stress in salt treated leaves (Figure 6 *A,B*) parallels previous observations that stress-induced GST gene expression peaks within 3 h of stress onset, followed by recovery to pre-induction levels by 12 h (126). Collectively, this suggests that glutathione metabolism represents an early-stage stress adaption, but not one that is sustained long-term.

### Agricultural implications

With global climate change continuing to deteriorate agricultural productivity through unpredictable environmental changes and grow seasons, it is necessary to generate crop varieties with higher tolerance to extreme conditions such as high salinity and drought. Further, with bioengineering / breeding of climate resilient crops requiring immediately actionable data, and proteins representing the effectors of the plant cell, next-generation quantitative proteomics workflows that can balance depth and speed of analysis are required. Correspondingly, we devised a quantitative proteomic workflow that fulfils this requirement and shows that quantitative proteomics can be readily utilized to generate time-resolved datasets relating to critical crop challenges. Our multi-dimensional proteomic dataset relating to root and shoot salt and osmotic stress responses across a 24 h time-course offers a comprehensive resource for future targeted gene / protein characterization as well as downstream breeders and agronomists evaluating existing populations for beneficial traits. For example, through the targeted characterization of just two clusters of up-regulated DAPs, we highlight the depth of our dataset and the utility of measuring time-related proteome changes under abiotic stress conditions.

## Summary

In this study, we successfully developed a new fast-flow, short-gradient multi-CV FAIMSpro BoxCarDIA acquisition method designed to capture time-resolved, proteome-wide protein abundance changes in plants. We demonstrate its utility by quantifying nearly 10,000 proteins from *Arabidopsis* seedling shoots and roots subjected to either salt or osmotic stress conditions. Our findings resolve a diversity of abiotic stress responses at the protein-level across these plant organs, providing novel, time-resolved insight into how plants respond to abiotic stress, in addition to critical new information for the breeding and/or biotechnological creation of climate resilient crops.

## Abbreviations

DDA: Data dependent acquisition
DIA: Data Independent acquisition
FAIMS: high field asymmetric waveform ion mobility spectrometry
ACN: acetonitrile
FTA: trifluoroacetic acid
FA: formic acid
LC: Liquid chromatography
*m/z*: mass-to-charge
MS/MS: tandem mass spectrometry
MS: mass spectrometry
PSMs: peptide spectrum matches

## Acknowledgements

The authors thank the Natural Sciences and Engineering Research Council of Canada and Canada Foundation for Innovation for funding this work. The authors also thank Jack Moore of the Alberta Proteomics and Mass Spectrometry Facility for assistance with mass spectrometer operation and maintenance.

## Supplemental Data

**Supplemental Data 1:** Experimental Meta Data

**Supplemental Data 2:** Complied Quantitative Proteome Data - Shoots

**Supplemental Data 3:** Complied Quantitative Proteome Data - Roots

**Supplemental Data 4:** Complied Quantitative Proteome Data – Shoot / Root – Abiotic Stress Overlap

**Supplemental Data 5:** GO enrichment – Time-Point Analysis

**Supplemental Data 6:** K-mean Cluster - Roots

**Supplemental Data 7:** K-mean Cluster - Shoots

**Supplemental Data 8:** K-mean Cluster - GO BP Enrichment - Root – Osmotic Stress

**Supplemental Data 9:** K-mean Cluster - GO BP Enrichment - Shoot – Salt Stress

